# Ganaxolone, an approved therapy for CDKL5-Deficiency Disorder, is an inhibitor of PTP1B

**DOI:** 10.1101/2025.10.31.685887

**Authors:** Imanol Zubiete-Franco, Qingting Hu, Nicholas K Tonks

## Abstract

CDKL5-deficiency disorder or CDD, which results in intellectual disability, speech and motor deficits, and seizures that can start as early as six weeks after birth, is caused by de novo mutations in the *CDKL5* gene. In early 2022, the FDA approved Ganaxolone (Commercial name: Ztalmy) for treatment of seizures in CDD patients 2 years and older. Ganaxolone has been reported to act as a GABAA receptor agonist that helps reduce neuronal excitability; however, based on its chemical structure we hypothesized that ganaxolone may also act as a PTP1B inhibitor. We observed that ganaxalone was able to inhibit PTP1B activity both in an in vitro assay of enzyme activity and in different cell models, in a comparable way to known inhibitors of PTP1B, including MSI-1436, which has a similar chemical structure. Additionally, inhibition of PTP1B in differentiating SH-SY5Y cells increased TRKB/BDNF signaling. This effect was prominent in CDKL5-KO cells, where inhibition of PTP1B brought TRKB levels and BDNF signaling to levels similar to those of wild-type cells and the observed signaling changes also coincided with restoration of cellular morphology. Finally, loss of CDKL5 function resulted in increased levels of PTP1B. Our results suggest that, like in Rett syndrome, targeting PTP1B may be a beneficial therapeutic strategy for CDD.

## Introduction

Rett-like syndromes, which include Rett syndrome, CDKL5-deficiency disorder (CDD) and FOXG1-syndrome, are rare yet devastating neurological disorders that share symptoms in common, including severe seizures and developmental delays, encompassing intellectual disability, autistic traits, speech and sleep deficits and motor deficits(1). Despite their similarities, these diseases are caused by distinct *de novo* mutations. Rett syndrome is caused by mutations in the methyl-CpG binding protein 2 (MECP2) transcription factor(2), CDD is caused by mutations in cyclin-dependent kinase like-5 (CDKL5), a protein Ser/Thr kinase(3, 4), and FOXG1-syndrome is caused by mutations in the forkhead box protein G1 (FOXG1) transcription factor(5). These mutations generally result in a reduction of brain size and weight and, at a cellular level, neurons display disruption of normal density and morphology(1). It is interesting to note that restoration of the mutated gene to wild type can repair some of the neuronal loss of function in different models(6–9) suggesting that these are not irreversible neurodegenerative conditions but, instead, the symptoms might be controlled with an appropriate treatment. Although gene therapy approaches may ultimately prove to be curative, the challenges of appropriate neuronal delivery and the dangers of over expressing the target protein are major obstacles at this time(10, 11). This has encouraged investigation into alternative therapeutic strategies to provide symptomatic relief, which may offer a quicker and safer approach to treatment in the short term.

These Rett-like syndromes feature aberrant neuronal development with reduced dendrite branching and spine density, improperly maintained synapses and deficient or shortened axons that result in abnormal phosphorylation-dependent signaling, culminating in an imbalance between inhibitory and excitatory activity in the brain(1, 12, 13). This suggests that, despite their distinct underlying causes, there may be common changes in signaling in these syndromes. Previously, our lab demonstrated that the loss of MECP2 leads to increased levels of the protein tyrosine phosphatase PTP1B(14), which is a crucial regulator of neuronal signaling pathways(15, 16). For example, PTP1B is known to dephosphorylate and inactivate the receptor tyrosine kinase TRKB(14, 17), which detects and transmits signals from the neurotrophin brain-derived neurotrophic-factor (BDNF). BDNF is important for the regulation of cell survival, synapse maintenance, and neuronal development(18). Both genetic ablation of *Ptpn1,* the gene encoding PTP1B, or inhibition with small-molecule-inhibitors, resulted in improved BDNF signaling and an improvement in symptoms in mouse models(14, 17). These data suggest that modifying signal transduction pathways with small-molecule drugs may offer a potential strategy for improvements to quality of life in Rett-like syndrome patients.

In the face of technical challenges to gene therapy, drug discovery efforts in CDD have focused on symptomatic relief. CDD patients suffer a high seizure burden, starting as early as three months of age, and often develop resistance to current anti-seizure medications(19–22). In 2022, the synthetic neurosteroid Ganaxolone, under the trade name ZTALMY(23), was approved by the FDA as a treatment for seizures in CDD patients two years and older(24–26). Ganaxolone has been reported to act as an agonist of the γ-aminobutyric acid-A (GABAA) receptor, resulting in a reduction in neuronal excitability and lowering the probability of suffering a seizure(24–26). The effects of treatment with ganaxolone were also reported to be maintained in over 85% of patients after up to two years of treatment(27). Although ganaxolone is considered an allosteric modulator of GABAA receptor subunits(28), there is also evidence that it may exert off-target effects by binding membrane progesterone receptors(29). In addition, we noticed that the structure of ganaxolone contains a similar steroid core to several aminosterol compounds that have been identified as allosteric inhibitors of PTP1B, including MSI-1436 and DPM-1001(30, 31). This suggested to us that inhibition of PTP1B activity may be one of the features underlying the mechanism of action of ganaxalone.

In this study, we report that PTP1B was inhibited by ganaxolone in assays of phosphatase activity in vitro, with similar potency to established allosteric inhibitors of the phosphatase. This focused our attention on testing the hypothesis that, as in Rett syndrome models, inhibition of PTP1B function may ameliorate some of the consequences of loss-of-function mutations in *Cdkl5* in CDD models. On this basis, we present data to show similar effects of inhibition of PTP1B function and ganaxalone treatment in a neuronal cell model. These data focus attention on possible new interpretations of the therapeutic effects of ganaxalone in CDD and the potential for therapeutic intervention at the level of the changes in signaling that underlie the disease.

## Results

### Ganaxolone is an inhibitor of PTP1B *in vitro*

Ganaxolone is a synthetic neurosteroid with a chemical structure similar to that of known aminosterol allosteric inhibitors of PTP1B, such as MSI-1436 and DPM-1001(30, 31) (Fig. 1A). Using DiFMUP (6,8-Difluoro-4-Methylumbelliferyl Phosphate) as a substrate, we tested ganaxolone as an inhibitor of PTP1B in an *in vitro* phosphatase assay, and compared its effects with MSI-1436 (Fig. 1B). Ganaxolone and MSI-1436 were able to inhibit PTP1B with similar potency (IC50: 760 nM vs. 660 nM respectively).

**Figure 1:**
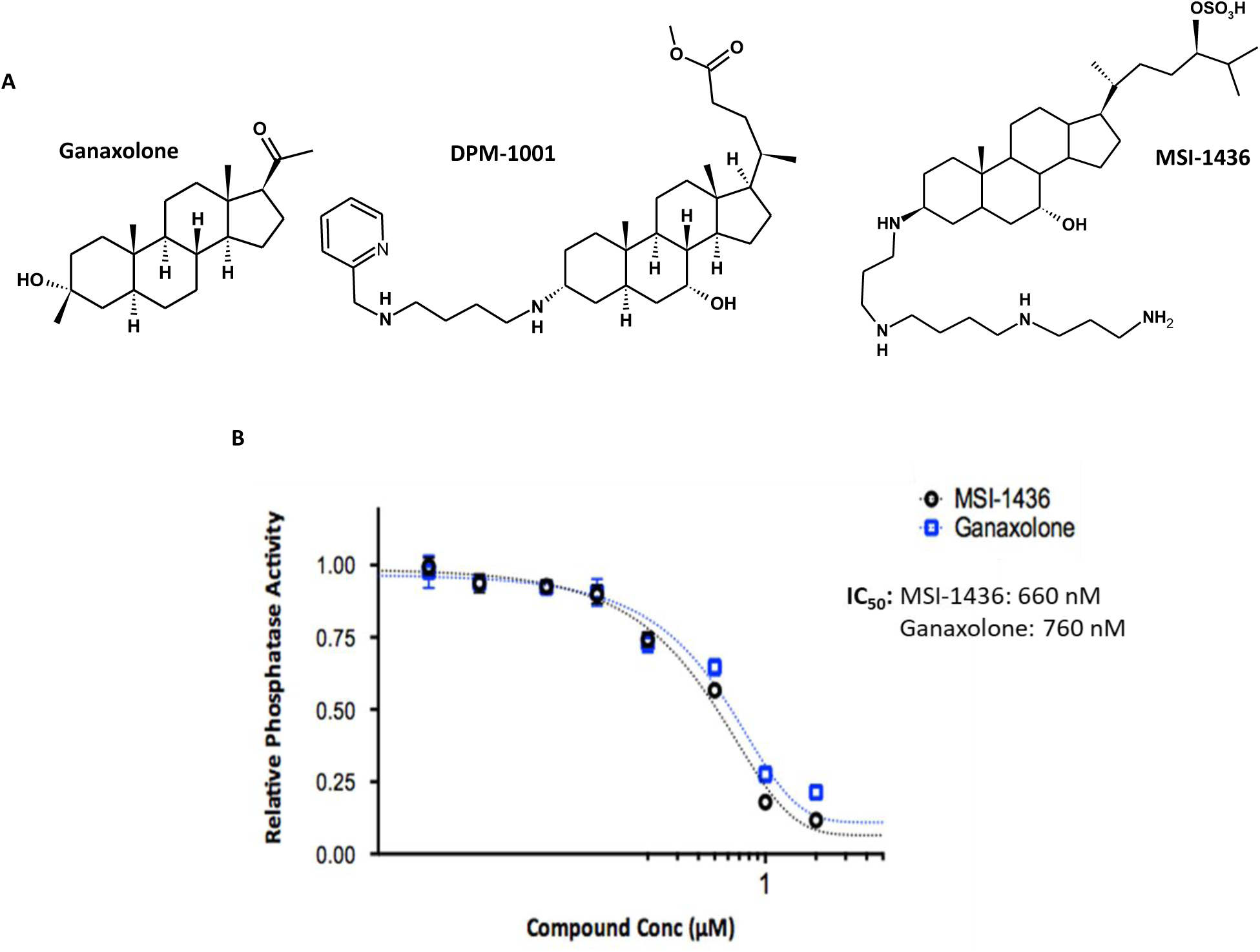
**Ganaxolone is a PTP1B inhibitor**. **A)** Comparison of the structures of ganaxolone, DPM-1001, and MSI-1436. **B)** Comparison of the inhibitory effects of ganaxolone and MSI-1436 in a PTP1B phosphatase activity assay using DiFMUP as a reporter.

To test whether these results could be replicated in a cellular model, we measured the effect of ganaxolone on tyrosine phosphorylation of the epithelial growth factor receptor (EGFR) in human 293T cells. We observed an increase in EGF-induced receptor phosphorylation after one hour of drug pre-treatment compared to vehicle-treated cells. This increase was comparable to that observed after treatment with MSI-1436 (Fig. 2A, Suppl. Fig. 1A) and suggests that ganaxolone can exert inhibitory effects on PTP1B activity in cells. The same experiment in a 293T-PTP1B-KO cell line resulted in similar EGFR phosphorylation levels in all conditions (Fig. 2A) suggesting that the effect we saw with ganaxolone was due to its ability to inhibit PTP1B.

**Figure 2:**
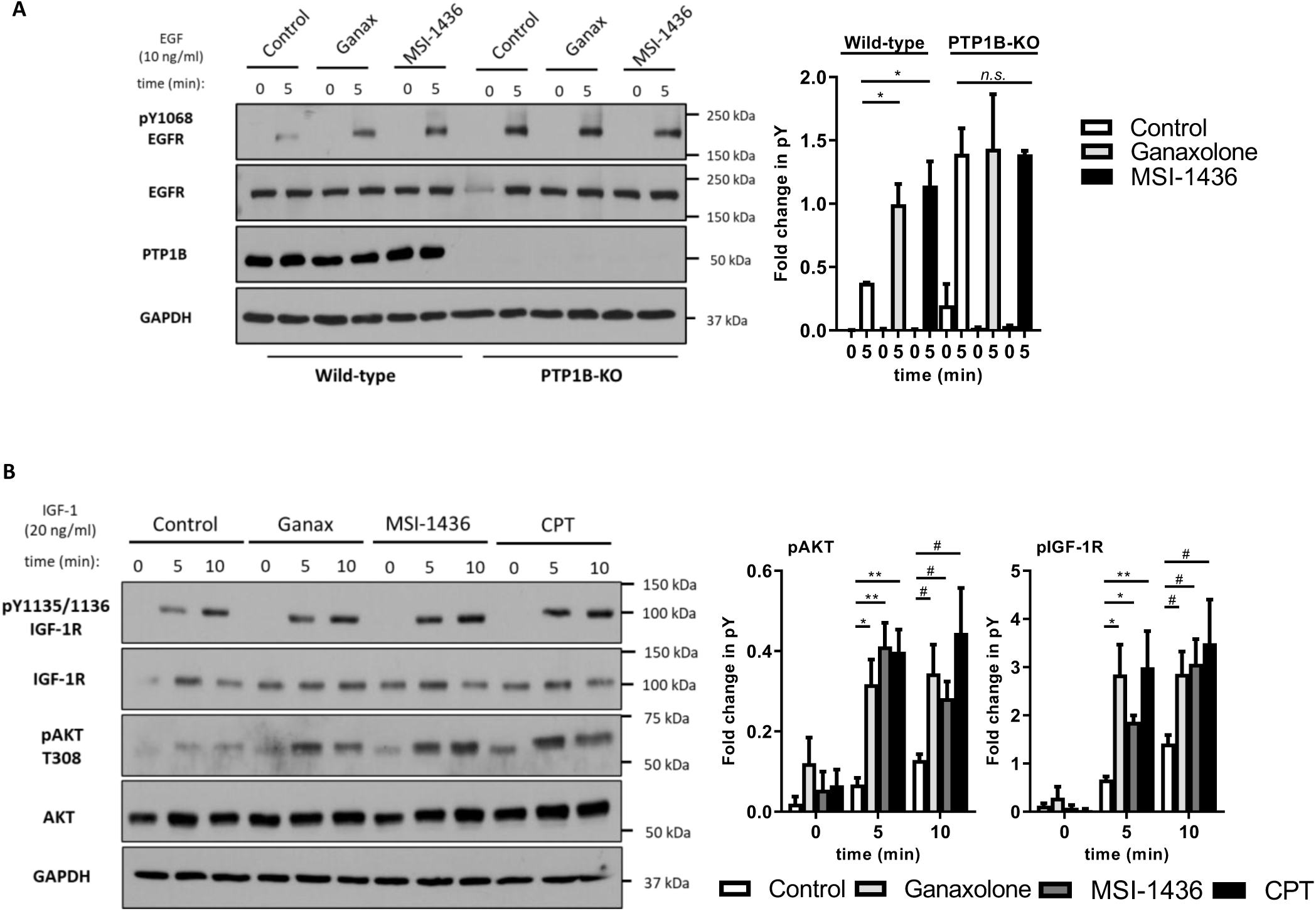
Ganaxolone inhibits PTP1B in cell models. **A)** Response of human 293T wild type cells and 293T PTP1B-KO cells to EGF (10 ng/ml, 5 minutes) after 1 hour pre-treatment with vehicle (Control), ganaxolone (5 μM, Ganax), or MSI-1436 (4 μM, MSI-1436). **B)** Response of undifferentiated SH-SY5Y cells to IGF-1 (20 ng/ml), including AKT activation, after 1 hour pre-treatment with vehicle (Control), ganaxolone (5 μM, Ganax), MSI-1436 (4 μM, MSI-1436), or CPT-157633 (4 µM, CPT) Blot quantitation presented to the right. */# = p<0.05, **/## = p<0.01, n.s. = not significant.

We repeated the experiments in the human neuroblastoma derived SH-SY5Y cells, which is a model used to study different neuronal processes(32). Treating undifferentiated SH-SY5Y cells for 10 minutes with IGF-1 after 1 hour of drug pre-treatment resulted in increased IGF-1 receptor (IGFR) tyrosine phosphorylation in a similar manner to that observed in 293T cells. Again, ganaxolone had a similar effect to MSI-1436 (Fig. 2B). Treatment with CPT-157633, a structurally distinct PTP1B inhibitor, also enhanced signaling, including downstream activation of AKT, in a comparable manner to ganaxalone (Fig. 2B, Suppl. Fig. 1B). These data are consistent with ganaxolone having an inhibitory effect on PTP1B function in a neuronal cell model.

### PTP1B inhibition increases TRKB expression and signaling during differentiation

The SH-SY5Y cell line can be differentiated into mature neuron-like cells that develop an axon and dendrites expressing the appropriate receptors(32). Treatment with retinoic acid (RA) over 6 days induces a morphological change from an epithelial-like appearance to branched neuronal-like structures that express neuronal markers(32). CDKL5-deficient neurons have reduced cell length, loss of dendrite branching, reduced cargo trafficking, and unstable synapses(12, 33, 34).

To test the impact of PTP1B inhibition on the differentiation of SH-SY5Y cells, we treated the cells with retinoic acid in the presence of ganaxolone, MSI-1436, or vehicle control, for six days (Fig. 3A). After six days of differentiation, it would be anticipated that SH-SY5Y cells would have detectable levels of TRKB and be able respond to BDNF(35). We detected an increase in receptor levels when cells were treated with both ganaxolone and MSI-1436 during differentiation when compared to control (Fig. 3B) suggesting that PTP1B might play a role in either receptor expression or transport.

**Figure 3:**
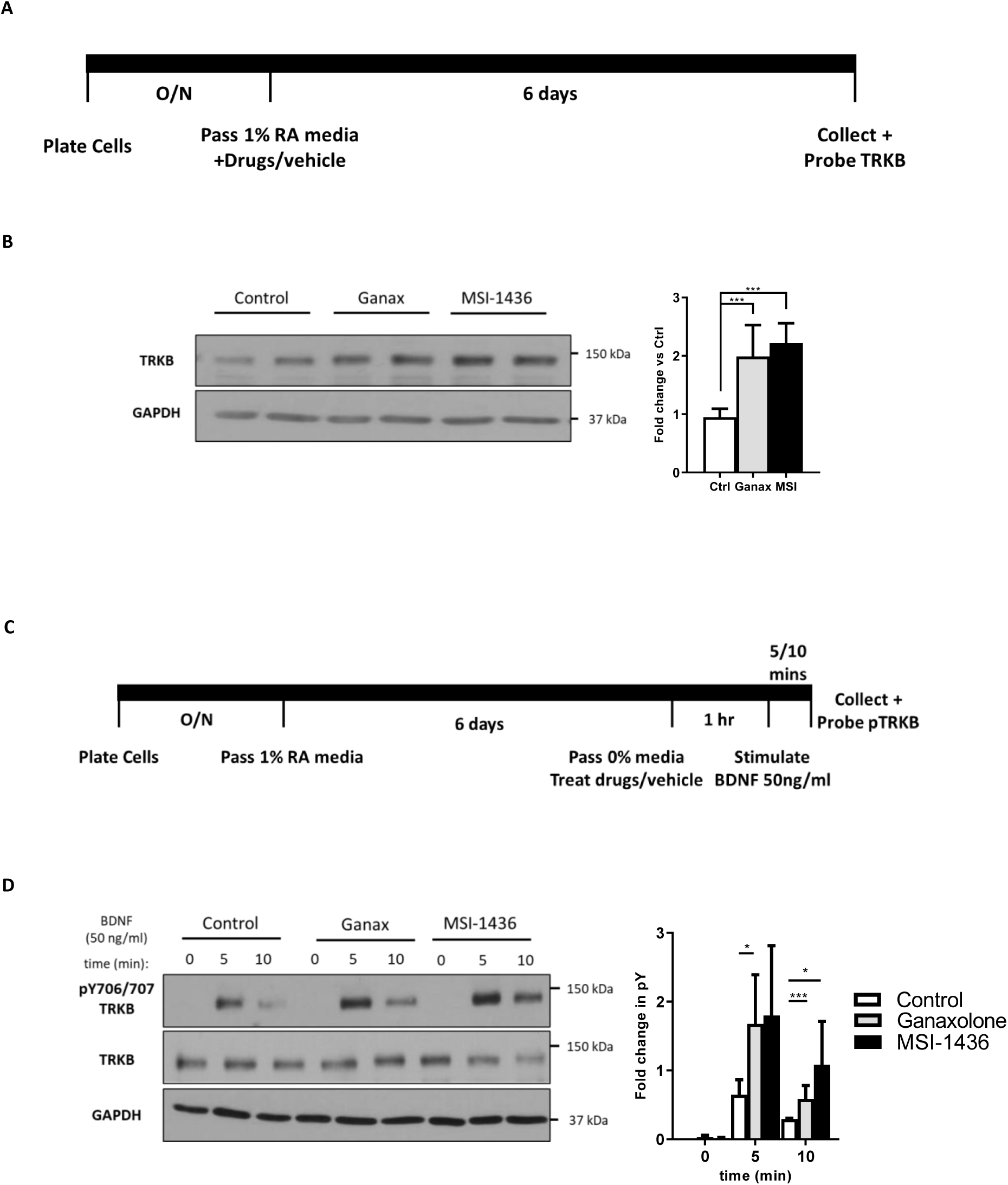
PTP1B inhibition increases TRKB receptor expression and signaling. **A)** Graphical depiction of the experimental procedure for figure 3B. **B)** 10 µM RA differentiating SH-SY5Y cells were treated with vehicle (Control/Ctrl), ganaxolone (5 µM, Ganax), or MSI-1436 (1 µM, MSI-1436) for 6 days. Cells were collected and probed for receptor levels. **C)** Graphical depiction of the experimental procedure for figure 3D. **D)** 6-day RA differentiated SH-SY5Y cells were pre-treated for 1 hour with vehicle (Control/Ctrl), ganaxolone (5 µM, Ganax), or MSI-1436 (4 µM, MSI-1436) and then stimulated with BDNF (50 ng/ml) for 5/10 minutes. Cells were collected and probed for receptor phosphorylation. Blot quantitation presented to the right. *** = p<0.001.

BDNF plays an important role in neuronal survival, development, and synapse maintenance(18). The changes in levels of TRKB that accompany differentiation would be expected to result in changes in downstream signaling. Previously, we had reported beneficial effects of inhibition of PTP1B in *Mecp2*-deficient mouse models of Rett syndrome that correlated with enhanced signaling through TRKB(14). Therefore, we examined potential links between PTP1B, CDKL5, and BDNF signaling in differentiated SH-SY5Y cells. We differentiated SH-SY5Y cells with retinoic acid, then treated them with vehicle, ganaxolone or MSI-1436 for one hour, and stimulated with BDNF for up to 10 minutes (Fig. 3C). Drug-treated cells displayed increased phosphorylation of TRKB after 5 minutes when compared to vehicle treated cells (Fig. 3D). Again, the effect of ganaxolone was similar to that of MSI-1436, further reinforcing the potential impact of ganaxalone on PTP1B function.

### Ablation of CDKL5 leads to elevated expression of PTP1B

Inactivating mutations in MeCP2 that underlie Rett syndrome are accompanied by changes in gene expression that alter the balance between inhibitory and excitatory neuronal circuits in the brain(13). MeCP2 influences the expression of several genes, of which *PTPN1* (encoding PTP1B) is one. Overall, PTP1B regulates signaling pathways of fundamental importance to brain function, including BDNF/TRKB and insulin signaling, providing a clear mechanistic basis for the observed beneficial effects of PTP1B inhibition in Rett models. With this in mind, we examined the effect of *CDKL5* knockout on expression of PTP1B. To mimic the effects of loss-of-function mutations in CDKL5, which would compromise its kinase function, we created a CDKL5-knockout SH-SY5Y cell line (C5KO) using the CRISPR-CAS9 method. As expected, phosphorylation of Ser222 in EB2 was decreased in C5KO cells (Fig 4A). Interestingly, as in the *Mecp2^-/-^* mouse model(14), we saw a dramatic increase in PTP1B levels when CDKL5 was ablated (Fig. 4A). This increase would be expected to suppress critical signaling pathways and further implicates PTP1B as a regulator of signaling events triggered by loss of CDKL5 function.

**Figure 4:**
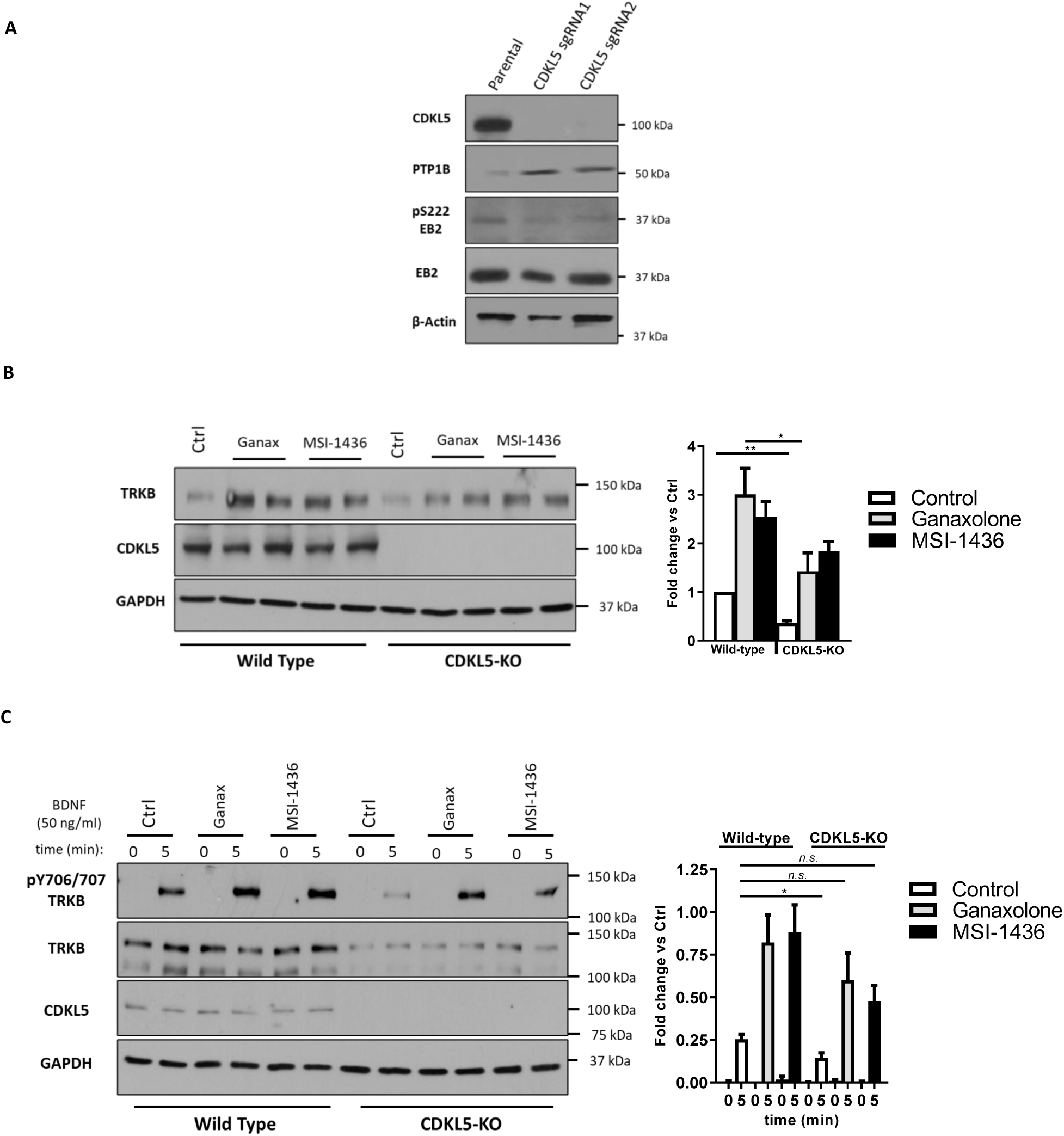
PTP1B inhibition can overcome CDKL5-KO signaling deficits. **A)** Characterization of CDKL5-KO SH-SY5Y (C5KO) cells. **B)** 10 µM RA differentiating SH-SY5Y wt and C5KO cells were treated with vehicle (Control/Ctrl), ganaxolone (5 µM, Ganax), or MSI-1436 (1 µM, MSI-1436) for 6 days. Cells were collected and probed for receptor levels. **C)** 6-day RA differentiated SH-SY5Y wt and C5KO cells were pre-treated for 1 hour with vehicle (Control/Ctrl), ganaxolone (5 µM, Ganax), or MSI-1436 (4 µM, MSI-1436) and then stimulated with BDNF (50 ng/ml) for 5 minutes. Cells were collected and probed for receptor phosphorylation. Blot quantitation presented to the right. * = p<0.05, ** = p<0.01.

To determine the impact of loss of CDKL5 function on signaling, we repeated the previous experiments using the C5KO cells (Fig. 3A and 3C). We differentiated C5KO cells with retinoic acid for six days, either in the presence of ganaxolone or vehicle control, and then checked cells for receptor levels. C5KO cells displayed reduced levels of receptor when compared to control; however, ganaxolone or MSI-1436 treatment resulted in a partial rescue of the levels of TRKB and bring it closer to wild type levels (Fig, 4B). Furthermore, as expected, BDNF signaling was decreased in vehicle-treated C5KO cells when compared to wild type; however, TRKB phosphorylation was increased close to wild-type levels in C5KO cells treated with ganaxolone or MSI-1436 for one hour (Fig. 4C).

### Loss of PTP1B rescues morphological defects associated with CDKL5 knockout in differentiated SH-SY5Y cells

In Rett-like syndromes, neurons display a variety of morphological abnormalities, including reduced size, shorter and less-branched dendrites, and decreased spine density(1). These changes highlight the deleterious effects on neuronal maturation and synapse function that underlie the disease. Using differentiated SH-SY5Y cells as a model, we observed that ablation of CDKL5 decreased the length of the neuron-like projections; in contrast, the length of the neuron-like projections was increased following ablation of PTP1B (Fig. 5A) Interestingly, the effect of CDKL5 loss was rescued by ablation of PTP1B in double-knockout cells (dKO) (Fig. 5A). Similarly, treatment of the CDKL5 knockout cells with ganaxalone, or PTP1B inhibitor, rescued the decrease in cell length caused by CDKL5 suppression alone (Suppl. Fig. 2).

**Figure 5:**
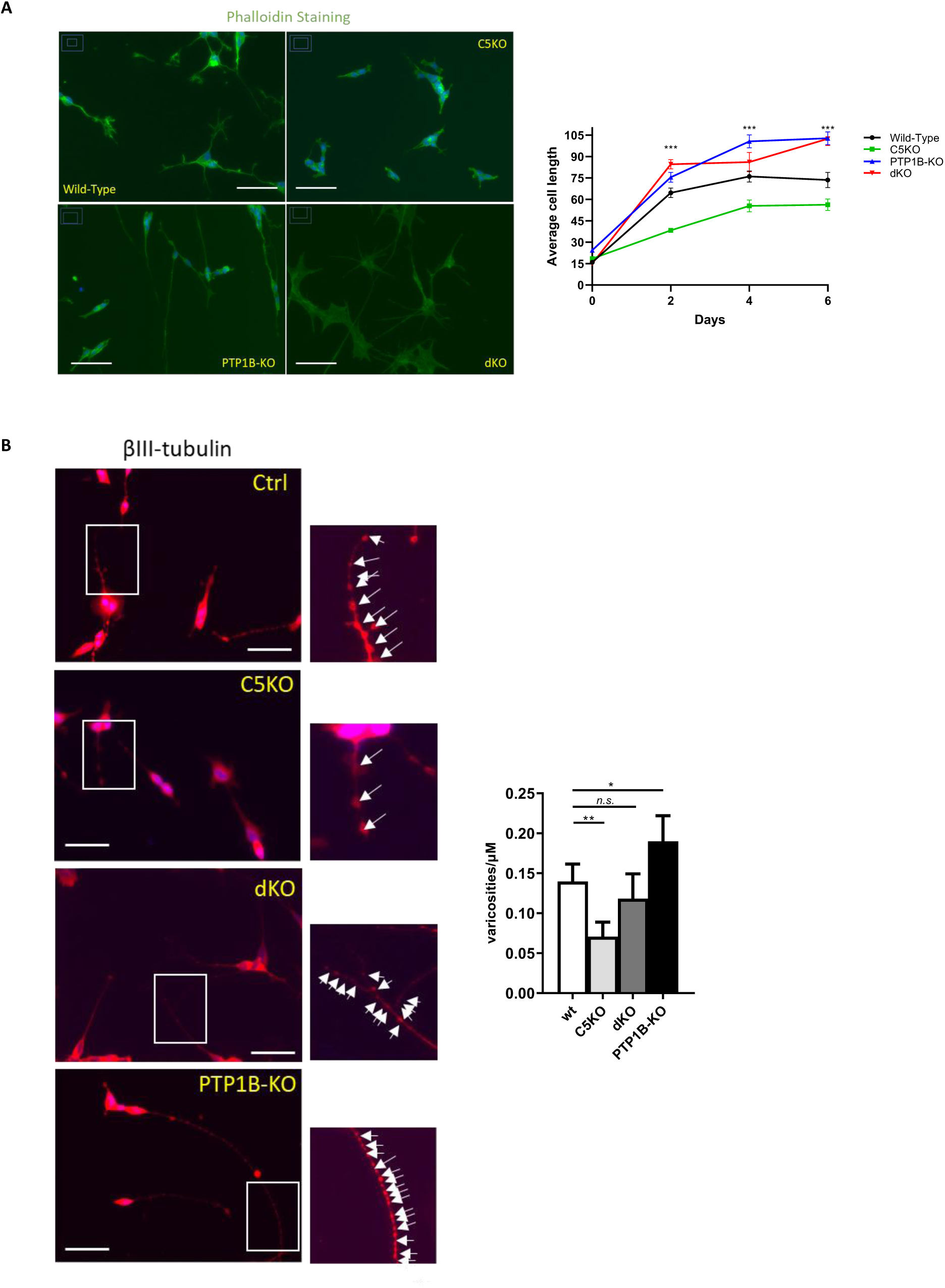
PTP1B inhibition impacts cellular morphology. **A)** 2-to-6-day RA differentiated SH-SY5Y cells (Wild type), CDKL5-KO cells (C5KO), PTP1B-KO cells and CDKL5/PTP1B doubleKO cells (dKO) were fixed at indicated timepoints and probed for phalloidin to measure cell length. Average cell length is displayed to the right. Pictures are representative of 6-day cultures. *** is a comparison between the C5KO and dKO cells. **B)** The same SH-SY5Y cell lines as in **A** were differentiated for six days, fixed and probed for βIII-tubulin to detect varicosities. Higher magnification shows the enlargement of a cell segment included in the white box. White arrows represent a detected varicosity. Scale bars = 130µM. Graphical representation of varicosities/µM is presented to the right. * = p<0.05, ** = p<0.01, *** = p<0.001, n.s. = not significant.

Differentiated SH-SY5Y cells also display swellings, or varicosities, along the length of the neurite(32). Although these are not functional dendritic spines, and do not contain post-synaptic density markers such as PSD95, they do share features of presynaptic structures, such as boutons en passant(32). We observed that following loss of CDKL5, the number of these varicosities was decreased relative to wild-type differentiated SH-SY5Y cells; in contrast, the number of varicosities was dramatically increased following suppression of PTP1B (Fig. 5B). The double knockout rescued the loss of varicosities caused by disruption of CDKL5 (Fig. 5B).

Taken together, these results indicate that loss of PTP1B in a loss-of-function-mutant CDKL5 background not only restores deficiencies in signaling but also can restore a normal cell morphology. These results are consistent with the model that at least some of the effects of ganaxolone may be exerted through inhibition of PTP1B and that, as in the case of Rett-syndrome(14), inhibition of PTP1B may be a viable strategy for therapeutic intervention in CDD.

## Discussion

CDKL5, Cyclin-Dependent Kinase-Like 5, is a member of the CMGC (Cyclin-dependent kinases, Mitogen-activated protein kinases, Glycogen synthase kinases, and CDK-like kinases) group of protein kinases(36). It is encoded by an X-linked gene, is highly expressed in brain, and plays an important role in several steps underlying normal brain development, including cell survival and proliferation, neuronal migration and synaptic development(1, 33, 34, 37). Loss-of-function mutations in *CDKL5* lead to the rare (∼1 in 40,000 live births) neurodevelopmental disorder, CDD (CDKL5-Deficiency Disorder)(4, 38, 39). Among its hallmarks, this disease features early-onset seizures, motor and speech impairment, intellectual disability, and autistic-like features(19, 40–43). Originally referred to as Atypical Rett syndrome, it shares overlapping phenotypes with Rett syndrome, which is caused by inactivating mutations in the X-linked gene *MECP2*, encoding the transcriptional regulator methyl CpG binding protein 2(2). Nevertheless, unlike Rett syndrome patients, whose symptoms typically appear from 6-18 months of age, CDD patients display epileptic encephalopathy and severe developmental delays from birth(44). By 3 months of age, patients may experience multiple different kinds of seizures that are often refractory to standard medications(44). In fact, up to 80% of patients can experience 10-20 seizure attacks every day, rendering it a debilitating disease(44). As CDD is caused by mutations in a single gene, *CDKL5*, there is hope that gene therapy approaches may prove to be curative; however, at this stage, such approaches are technically challenging and remain in preclinical development(45). Instead, thus far, therapeutic strategies for CDD have focused on symptomatic relief, but with rather disappointing results overall(21, 22, 43, 46). In particular, current therapies have addressed seizures as the core feature of the disease and patients take multiple anti-seizure medications simultaneously.

The anti-seizure medication Ganaxolone (commercial name Ztalmy) is the first to be approved by the FDA for treatment of seizures in CDD(24–26). Ganaxalone is a neurosteroid, a synthetic analogue of allopregnanolone, and is reported to exert its anti-seizure effects by acting as a positive allosteric modulator of GABAA receptors(24–26). Ganaxalone has been reported to prevent seizures in rodent models(47). It displayed acceptable safety and tolerability, and reduced seizure frequency in an open-label phase 2 trial(48). In a double-blind, randomized, placebo-controlled, phase 3 clinical trial ganaxalone treatment resulted in a modest ∼30% decrease in seizure frequency compared to ∼7% in the placebo group(27, 49–51). Adverse effects associated with treatment occurred in 86% of the ganaxalone group (12% classified as serious) and 88% of placebo(27, 49–51). Overall, it was concluded that ganaxalone was useful as an adjunct therapy with existing anti-epileptic drugs, particularly in drug-resistant states, that its effects persisted over a two-year study, and that it was relatively well tolerated. Nevertheless, the doses of ganaxalone required to elicit effects in CDD patients are high (∼1.8g/day)(27, 50) and impacts on non-seizure outcomes were less clear(52). This suggests that it may be prudent to reassess whether the impact on GABAA receptor function explain all of the effects of ganaxalone in CDD. Perhaps alternative targets and mechanisms of action should be considered to achieve a broad impact on symptoms in the treatment of CDD.

As a protein kinase, CDKL5 has potential to regulate signal transduction and it would be expected that loss-of-function mutations in *CDKL5* would be manifested as changes in critical signaling events in the brain that support neuronal function. In fact, identification of the targets of CDKL5 could suggest deficiencies that underlie the etiology of CDD and suggest potential new druggable targets(12, 33, 34, 44). As in Rett syndrome, the development of mouse models has provided an important step forward in understanding the pathophysiology of the disease and the roles of CDKL5 in neuronal function and development(41, 53–57). These models display many key symptoms, including motor dysfunction, impaired learning and autistic-like features, while also manifesting critical abnormalities in neuronal structure and function(58–60). One problem has been that such models tend not to display spontaneous seizures, limiting their utility; however, more recent studies have now revealed spontaneous seizures in several CDD models(58, 60). Considering the insights gained from these animal models, we propose that PTP1B may represent a new, druggable target for CDD.

PTP1B regulates multiple pathways that are of fundamental importance to normal homeostasis. For example, PTP1B has been recognized as a negative regulator of insulin and leptin signaling pathways and as a highly validated therapeutic target for new approaches to treatment of diabetes and obesity(61). Also, PTP1B acts to suppress BDNF signaling through dephosphorylation of its receptor TRKB(14, 17); consequently, inhibitors of PTP1B would be expected to promote BDNF/TRKB signaling. Most significantly, we have validated PTP1B as a therapeutic target for Rett syndrome(14). Some Rett mouse models display obesity and leptin resistance, with insulin resistance also noted in some Rett patients, which suggested to us that PTP1B function might be altered in Rett syndrome. Previously, we demonstrated that the *PTPN1* gene, which encodes PTP1B, is a direct target of MECP2, which interacts with the *PTPN1* promoter and suppresses promoter activity. Furthermore, disruption of MECP2 function was associated with increased levels of PTP1B in RTT models. As expected, pharmacological inhibition of PTP1B, with multiple structurally and mechanistically distinct small molecule inhibitors, reduced the extent of glucose intolerance and improved metabolic status in the Rett syndrome mouse model. Of particular importance, the PTP1B inhibitors ameliorated a wide array of the effects of MECP2 disruption in Rett mice, including movement, behavior and heart function(14). Furthermore, we demonstrated that the elevated levels of PTP1B in Rett syndrome represent a barrier to BDNF signaling; inhibition of PTP1B led to increased tyrosine phosphorylation of TRKB in the brain, which augmented the response to BDNF(14). These observations suggest that PTP1B adopts a dominant role as an inhibitor of signaling in Rett syndrome, supporting inhibition of PTP1B function as a new, mechanism-based approach to treatment. Analysis of the phenotype of CDD mouse models suggests the possibility that inhibition of PTP1B may counter the impact of loss of CDKL5 function on neuronal development.

CDKL5 serves a pro-survival function in immature neurons and is required for normal dendrite formation. In CDKL5-knockout mice, there is increased neuronal precursor apoptotic cell death and decreased granule cell number, loss of dendrite development, and impaired hippocampus-dependent memory(37); these effects have been linked to impaired AKT signaling(37). Considering the role of PTP1B as a suppressor of PI3K-AKT signaling downstream of the insulin receptor(61, 62), one might expect that inhibition of PTP1B would re-establish AKT signaling and address this deficit in neuronal development. Similarly, a reduction in spine density has been reported in CDKL5-mutant mice, leading to impaired LTP; administration of exogenous IGF-1 rescued the defects in spine density and PSD95 expression(63). We have reported that PTP1B can suppress IGF-1 signaling(64), consequently, inhibition of PTP1B may also reverse this dendritic hypotrophy. In the perirhinal cortex (PRC), loss of CDKL5 led to reduction of dendrite length, spine density, and maturation, which could be ameliorated with TRKB agonists(65). Considering the ability of PTP1B to suppress TRKB signaling, these effects of loss of CDKL5 on neuronal development and function could be ameliorated by inhibition of PTP1B to promote BDNF signaling and neuronal health. Increased microglial activation has been reported in CDD mouse models, suggesting a neuroinflammatory component to the disease pathophysiology(66).

Treatment with LUTEOLIN, to inhibit neuroinflammation, ameliorates symptoms and improves hippocampal neurogenesis and dendritic maturation(67). PTP1B has been reported to promote microglia-mediated neuroinflammation(68) and we have demonstrated that PTP1B modulates microglial-dependent activation in a SYK-dependent manner(69), suggesting that inhibition of PTP1B may also reverse this inflammatory state. Finally, links are being established between brain metabolism and epilepsy(70), including the application of GLP-1 receptor agonists to reduce seizures(71, 72). Considering the central role of PTP1B in metabolic regulation, this may reflect another mechanism by which a specific PTP1B inhibitor may ameliorate the pathophysiology of CDD by modulating a wider range of signaling deficits.

In this study, we draw these aspects together with the demonstration that ganaxalone is a PTP1B inhibitor both in phosphatase activity assays in vitro and in cell-based models. Furthermore, its impact in cell models was mimicked by established allosteric inhibitors of PTP1B that are structurally related to ganaxalone. Finally, we demonstrate that, as in Rett syndrome(14), loss of CDKL5 results in higher levels of PTP1B, establishing an additional link between the phosphatase and the CDD phenotype. An interesting consequence of this effect is that it not only implicates PTP1B as a potential regulator of critical signaling events triggered by loss of CDKL5 function, but also the increased levels of the phosphatase may help to distinguish disease from normal cells and provide a greater range of doses over which the PTP1B inhibitors may exert their effects in a CDKL5-impaired cell background. It is interesting to note that overexpression of PTP1B in forebrain has also been linked to neuroinflammation, synaptic dysfunction, and cognitive impairment in obesity(73). Our data suggest that inhibition of PTP1B by ganaxalone may reflect a crucial “off-target” effect of the drug that may contribute to some of its impact on CDD. Although designed as a GABAA receptor agonist, considering links between metabolic dysfunction, neuroinflammation, and seizure disorders, it is possible that inhibition of PTP1B by ganaxalone may contribute to its anti-seizure effects. Furthermore, our data suggest that by optimizing ganaxalone for PTP1B inhibition, or using another PTP1B-specific inhibitor, we may open up additional effects, including in both neurons and glia, that may address a broader range of symptoms of CDD and thus provide a more effective therapy.

## Materials and Methods

### Reagents

All common reagents were obtained from ThermoFisher Scientific (Waltham, MA) or Sigma-Aldrich (St. Louis, MO). Ganaxolone (G7795) and human Insulin (91077C) were obtained from Sigma-Aldrich. Human endothelial growth factor (#AF-100-15) and human Insulin-like growth factor 1 (#100-11) were obtained from Peprotech (Waltham, MA). Retinoic Acid (#0695) was from Tocris Bioscience (Bristol, UK).

### Antibodies

Total TRKB (#4603), pY706/707 TRKB (#4621), total EGFR (#4267), pY1068 EGFR (#3777), total IGF-1R (#9750), pY1135/1136 IGF-1R (#3024), Total Insulin receptor (#3025), pY1150/1151 Insulin receptor (#3024), Total Akt (#9272), and pT308 Akt (#9275) were obtained from Cell Signaling Technology (Danvers, MA). Total EB2 (mab71717) and pS222 EB2 (pab01032-P) were from Covalab (Bron, France). CDKL5 (#703756) was from ThermoFisher Scientific. PTP1B (HPA012542) and β-III Tubulin (T2200) were from Sigma-Aldrich. GAPDH (ab125247), β-Actin (ab6276), phalloidin (ab176753) and NeuN (ab279297) were from Abcam (Waltham, MA). Secondary anti-rabbit HRP (ab97501) and anti-mouse HRP (ab97023) antibodies were from Abcam.

### Purification of wild-type PTP1B

Wild-type PTP1B containing a His-tag was expressed in Escherichia coli BL21 (DE3)-RIL (ThermoFisher Scientific) using LB broth. Cells were resuspended in lysis buffer (50 mM HEPES pH 7.2, 150 mM NaCl, 10 mM imidazole, 2 mM TCEP) with cOmplete EDTA free Protease Inhibitor Cocktail (Roche), and then lysed using a sonicator at 4°C. Lysates were clarified by centrifugation. Initial protein purification was by gravity flow, nickel column chromatography. Proteins were used immediately or stored at -80°C in in 50 mM HEPES pH 7.4, 100 mM NaCl, 2 mM dithiothreitol and 25 % glycerol.

### PTP1B phosphatase assay

PTP assays were performed in black polystyrol 96-well plates (Corning) using Difluoro-4-methylumbelliferyl phosphate (DiFMUP, Invitrogen, #D22065) as a substrate. DiFMUP (10 μM) was added to assay buffer (50 mM HEPES, 100 mM NaCl, 0.1% BSA, 2 mM DTT, 2 mM EDTA, pH 6.5) containing 5 nM purified PTP1B and ganaxolone or MSI-1436 (0-5 µM) in a final volume of 100 μl. The fluorescence emitted at 450 nm was monitored continuously for 240 min using a Gemini XPS fluorescence plate reader (Molecular Devices, San Jose, CA).

### Cell culture

293T (ATCC, Manassas, VA) and 293T-PTP1BKO cells were maintained in DMEM (Gibco, Waltham, MA) 10% FBS (Corning, NY) and 1% penn/strep (Gibco). To test the EGF response, 160,000 cells/well were serum starved overnight and then treated with vehicle/ ganaxolone (5 µM) or MSI-1436 (4 µM) for 1 hour. Treated cells were then stimulated with EGF (20 ng/ml or 10 ng/ml (PTP1BKO)) for 5 and 10 minutes and collected. SH-SY5Y cells (Cytion, Germany) were maintained in DMEM:F12 (Gibco) with 10% FBS and 1% penn/strep. To test IGF-1 or insulin response, 200,000 cells/well were serum starved and treated with vehicle/ ganaxolone (5-20 µM), MSI-1436 (4 µM) or CPT-157633 (4 µM) for 1 hour. Treated cells were then stimulated with IGF-1 (20 ng/ml) or insulin (10 nM) for 5 to 10 mins and collected. All cells were lysed and collected in RIPA buffer (25 mM HEPES pH 7.5, 150 mM NaCl, 0.25 % Deoxychloate, 10 % Glycerol, 25 mM NaF, 10 mM MgCl2, 1 mM EDTA, 1 % Triton X-100, 0.5 mM PMSF, 10 mM Benzamidine, cOmplete protease inhibitor cocktail). Soluble proteins were harvested by centrifugation and quantitated using the BCA method. Total proteins in the cell lysates were separated by SDS-PAGE and the specific proteins were monitored with their corresponding antibodies. GAPDH or β-Actin were used as loading controls. Blot images were quantified using ImageJ software.

### SH-SY5Y differentiation

SH-SY5Y cells were differentiated by culturing 160,000 cells/well in DMEM:F12 with 1% FBS, 1% PS, and 10 µM Retinoic acid for 6 days. To determine the effects of PTP1B inhibition on receptor expression, differentiating cells were additionally cultured with vehicle, ganaxolone (5 µM) or MSI-1436 (1µM) and were collected at day 6. To determine TRKB phosphorylation levels, differentiated cells were treated with vehicle, ganaxolone (5 µM), or MSI-1436 (2 µM) for 1 hour. Treated cells were then stimulated with 50 ng/ml BDNF (#78005, Stemcell Technologies, Vancouver, Canda) for 5 to 10 mins and then collected.

### CRISPR-CAS9 Knockouts

The 293T-PTP1BKO was created by using the CRSIPR plasmid pDG459. pDG459 was a gift from Paul Thomas (Addgene plasmid #100901; RRID: Addgene_100901). Used sgRNAs are available in supplemental Table 1. Edited cells were selected by culturing the transfected cells with puromycin (3 µg/ml, GIBCO) for three days. Single cells were then selected by array dilution and probed for loss of PTP1B.

SH-SY5Y CRISPR-CAS9 knockout cells were created by lentiviral delivery of the CRISPR plasmid lentiCRISPR v2. lentiCRISPR v2 was a gift from Feng Zhang (Addgene plasmid #52961; RRID: Addgene_52961). Used sgRNAs are available in supplemental Table 1. Virus was created by co-transfecting 293T cells with plasmids lentiCRISPRv2 and packaging plasmids VSVG and psPAX (Gifts from Dr. Vakoc lab) using lipofectamine 3000 (ThermoFisher Scientific) and following the manufacturer’s instructions. Media was refreshed and virus present in media was collected at 24 and 48 hours after transfection. Viral media was centrifuged and then filtered through a 45 µm filter. SH-SY5Y cells were transformed by culture with 1 ml of the appropriate virus and polybrene (8 µg/ml) for 1 day. Media was then refreshed for another 24 hours and puromycin (1.5 µg/ml) was added for two to three days to select for edited cells. Cells were probed for loss of the specific gene.

### Immunofluorescence

SH-SY5Y cells (wt/C5KO/PTP1B-KO/dKO) were grown on coverslips and differentiated for six days using 10 μM RA in 1% DMEM:F12 medium. Cells were fixed in PBS 4% paraformaldehyde. Coverslips were then blocked in 3% FBS with 0.1% Triton-X for 1 hour at room temperature, followed by overnight incubation at 4°C with primary antibodies (Phalloidin, βIII-Tubulin or NeuN). After incubation, the coverslips were washed three times with PBS and incubated with fluorescent secondary antibodies for 2 hours at room temperature, along with DAPI (1:10000) to stain the nuclei, followed by two washes with PBS. Coverslips were mounted on slides with Ibidi mounting medium (Ibidi, Germany). Images were captured using either an Echo fluorescence microscope (Echo, San Diego, CA) or a Zeiss LSM 780 confocal microscope (Zeiss, Germany). Images were taken randomly. Images were analyzed using ImageJ software.

### Statistics

All data are presented as mean ± SEM and represent a minimum of three replicates. Significance was determined by using unpaired two-tailed Student’s T-test with a p<0.05 considered as significant.

## Acknowledgments. Funding

N.K.T. is the Caryl Boies Professor of Cancer Research at Cold Spring Harbor Laboratory. Research in the Tonks lab was supported by NIH grant R01CA53840, the CSHL Cancer Centre Support Grant CA45508, a grant from CART (Coins for Alzheimers’ Research Trust), and the Hansen Foundation.

## Abbreviations

The following abbreviations are used: CDD, CDKL5-deficiency disorder; CDKL5, Cyclin-dependent kinase like-5; MeCP2, methyl-CpG binding protein 2; FOXG1, forkhead box protein G1; BDNF, brain-derived neurotrophic-factor; PTP1B, protein tyrosine phosphatase 1B; GABAA, γ-aminobutyric acid-A; DiFMUP, 6,8-Difluoro-4-Methylumbelliferyl phosphate; EGFR, epithelial growth factor receptor; IGFR, Insulin-like growth factor 1 receptor; RA, retinoic acid; TRKB, Tropomysin receptor kinase B; C5KO; CDKL-5-knock out SH-SY5Y cells; dKO, CDKL-5/PTP1B-double knock out SH-SY5Y cells; CMGC, cyclin-dependent kinases, Mitogen-activated protein kinases, Glycogen synthase kinases and CDK-like protein kinases.

## Supporting Information

This article contains supporting information

## Data Availability

All data generated or analyzed during this study are included in this article.

