## Supplemental material for "Ganaxolone, an approved therapy for CDKL5-Deficiency Disorder, is an inhibitor of PTP1B"

### Supplemental Figure 1

A

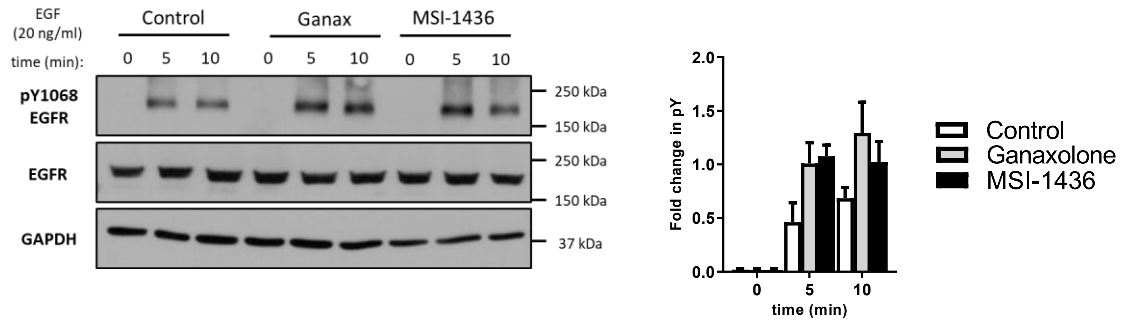


B

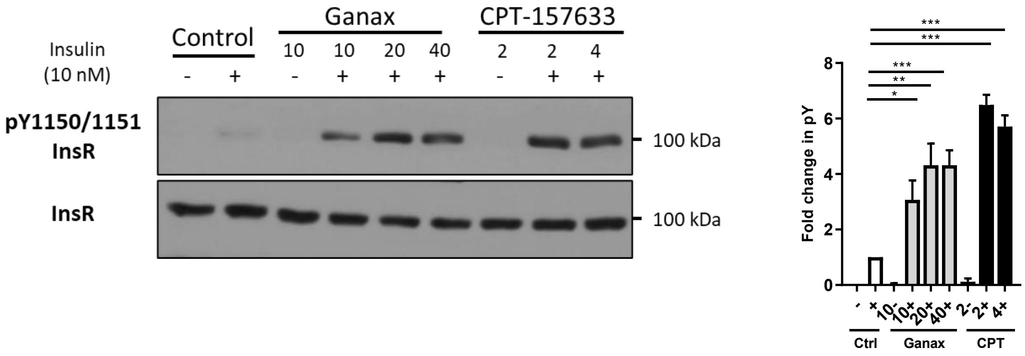


**Supplemental Figure 1: A)** Response of human 293T cells to EGF (20 ng/ml, 5/10 minutes) after 1 hour pre-treatment with vehicle (Control), ganaxolone (5 μM, Ganax), or MSI-1436 (4 μM, MSI-1436). **B)** Undifferentiated SH-SY5Y cells were pre-treated with drug (ganaxolone: 10-40 μM, CPT-157633: 2-4 μM) for 1 hour. Treated cells were then stimulated with 10 nM Insulin for 10 minutes, collected and probed for Insulin receptor phosphorylation. Immunoblot quantitation is shown on right. Ganax = ganaxolone, CPT = CPT-157633.

__Supplemental Figure 2__

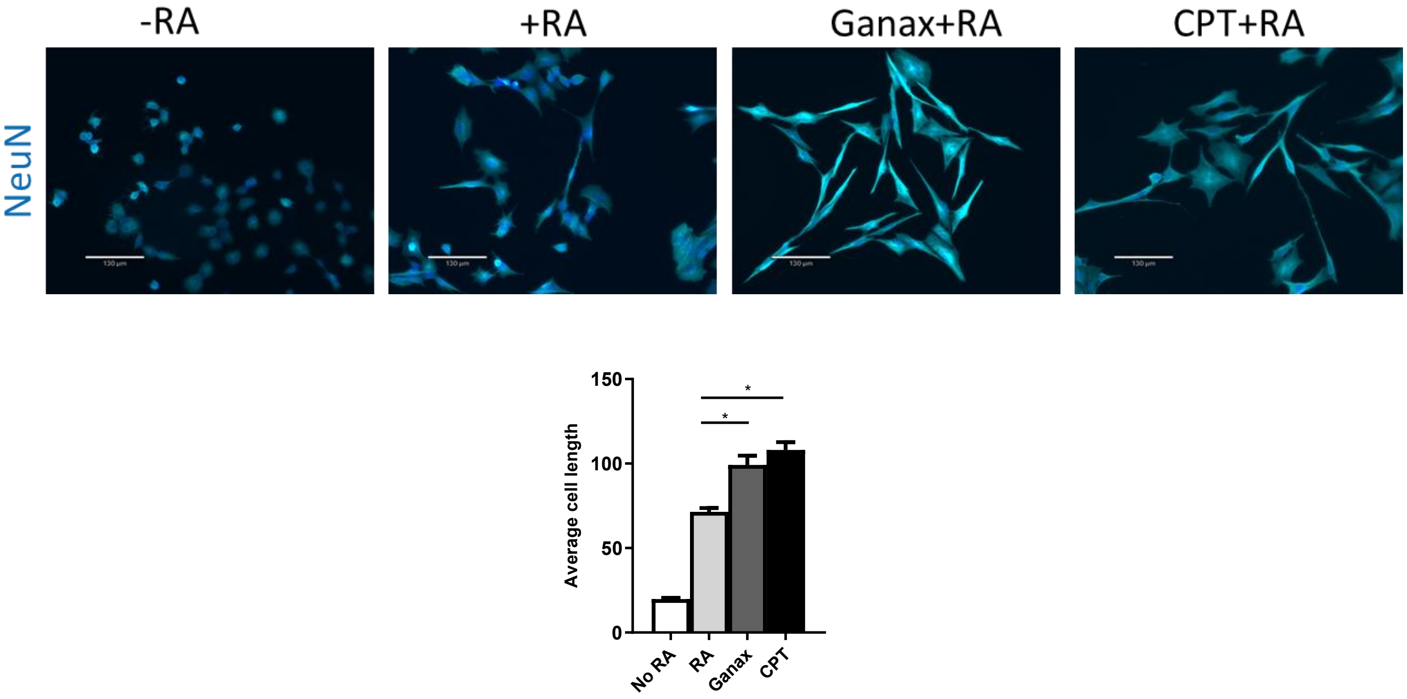


**Supplemental Figure 2:** Ganaxolone changes cell morphology in CDKL5-KO cells. C5KO cells were differentiated with RA for 3 days and then treated with vehicle or drugs for a further 3 days. Cells were then fixed and probed for NeuN. Average cell length for each condition displayed below. * = $p<0.05$. White line represents 130 μM. Ganax = ganaxolone (10 μM). CPT = CPT-157633 (4 μM).

### <u>Supplemental Materials and Methods</u>

**<u>Supplemental Table 1</u>:** List of sgRNAs used in the study.

| GENE | SEQUENCE (5'-3') |
|---|---|
| PTPN1 (293T) | **A: CAGTGACTTCCCATGTAGAG**<br>**CTCTACATGGGAAGTCACTG**<br><br>**B: GACGTCTCTGTACCTATTT**<br>**AAATAGGTACAGAGACGT** |
| CDKL5 (SH-SY5Y) | **sg1: TACGAGAGCTTAAAATGCTT**<br>**AAGCATTTTAAGCTCTCGTA**<br><br>**sg2: AATGCTTCGGACTCTCAAGC**<br>**GCTTGAGAGTCCGAAGCATT** |
| PTPN1 (SH-SY5Y) | **GTCTTTCAGTTGACCATAGT**<br>**ACTATGGTCAACTGAAAGAC** |

Sequences of the sgRNAs used in this study. The plasmid (pDG459) used to create PTP1B-KOs in 293Ts incorporates two sgRNAs (A and B) within the same plasmid. The two sgRNAs used in to create CDKL5-KO in SH-SY5Y were used to create two different clones.
